# Figure-ground organization in visual cortex for natural scenes

**DOI:** 10.1101/053488

**Authors:** Jonathan R. Williford, Rüdiger von der Heydt

## Abstract

Figure-ground organization and border-ownership assignment are essential for understanding natural scenes. It has been shown that many neurons in the macaque visual cortex signal border-ownership in displays of simple geometric shapes such as squares, but how well these neurons resolve border-ownership in natural scenes is not known. We studied area V2 neurons in behaving macaques with static images of complex natural scenes. We found that about half of the neurons were border-ownership selective for contours in natural scenes and this selectivity originated from the image context. The border-ownership signals emerged within 70 ms after stimulus onset, only ^~^30 ms after response onset. A substantial fraction of neurons were highly consistent across scenes. Thus, the cortical mechanisms of figure-ground organization are fast and efficient even in images of complex natural scenes. Understanding how the brain performs this task so fast remains a challenge.

**Significance Statement:** Here we show, for the first time, that neurons in primate visual area V2 signal border-ownership for objects in complex natural scenes. Surprisingly, these signals appear as early as the border-ownership signals for simple figure displays. In fact, they emerge well before object selective activity appears in infero-temporal cortex, which rules out feedback from that region as an explanation. Thus, “objectness” is detected by extremely fast mechanisms that do not depend on feedback from the known object-recognition centers.

## Introduction

Many visual tasks depend fundamentally on our visual system being able to organize low-level image features together as objects. This is a challenging problem for images of natural scenes in which object contours are buried in luminance and color variations produced by surface structure, illumination and shadows, and features from different objects are cluttered because of interposition in space. How the visual system is able to accomplish this is a major unresolved question of visual neuroscience. Border-ownership coding, discovered by Zhou et al. (2000), is an early neural correlate of this perceptual organization (see Williford and von der Heydt, 2013 for a review). Zhou et al. found that when an edge of a figure, such as a square, is aligned to the classical receptive field (CRF) of neurons in macaque visual cortex, some neurons will fire at a higher rate when the figure appears on one side of the CRF compared to the opposing side, even when the stimulus is locally ambiguous about the side of the figure. The underlying figure definition mechanisms may provide the structure for object-based attention (Qiu et al., 2007). However, border-ownership coding has not yet been demonstrated for natural scenes. The same is true for other neural correlates of figure-ground organization such as the enhancement of activity over figure regions (Lamme, 1995; Zipser et al., 1996; Lee et al., 1998; Marcus and Van Essen, 2002).

The goal of the present study was to close this gap of knowledge. We did not know what result to expect. On the one hand, images of natural scenes are rich in information that might be used for figure-ground definition. On the other hand, these images are infinitely more complex than displays of simple geometrical figures. Models of border-ownership coding were mostly designed for geometrical figures (Zhaoping, 2005; Sakai and Nishimura, 2006; Craft et al., 2007) and when such models were tested on complex natural scenes their performance was modest (Sakai et al., 2012; Russell et al., 2014).

Since primate visual systems have no difficulty in segregating figure and ground in complex scenes it is commonly assumed that this is based on object knowledge. Thus, it is often assumed that border-ownership signals for complex natural scenes would depend on prior shape processing. Since shape selective neurons in inferotemporal cortex start to respond only about 80ms after stimulus onset and become shape selective even later (Brincat and Connor, 2004), this hypothesis predicts that border-ownership signals for natural scenes should be delayed compared to those for simple geometrical figures, which appear around 60-70ms (Sugihara et al., 2011).

We report here that a subset of neurons in visual area V2 are selective for border-ownership in images of natural scenes. When the CRF of such a neuron aligns with an occluding contour, the neuron will fire at a higher rate on average when the occluding object is located on the neuron’s preferred border-ownership side. Some neurons signal border-ownership consistently across different objects and scenes. The border-ownership signals are mainly driven by the image context, while an image patch covering the CRF produces only a small transient signal. Surprisingly, border-ownership signals for natural scenes emerge as early as 70ms after stimulus onset, which contradicts the traditional view which attributes image understanding to shape processing at levels much higher in the hierarchy.

## Results

We studied neurons in the visual cortices from 5 hemispheres of 3 male rhesus macaques which we will refer to as FR, BE and GR. Once spikes of a cell were isolated, its orientation and color selectivity was determined and its CRF was mapped with bars or contrast edges. Next, border-ownership selectivity was measured by placing an edge of a square across the CRF center with the square located either on one side or the other of the edge. Responses to both sides were recorded with either contrast polarity (flipping the colors of square and background), and this was done for two sizes of squares (3 and 8° visual angle on a side). Flipping side of square and contrast polarity produced pairs of configurations in which the figure was on opposite sides while the edge in the CRF was identical. The border-ownership signal was defined as the response difference associated with the location of the figure. The difference averaged over the two sizes of squares we call the border-ownership signal for the “standard square test”. Since it compares pairs of conditions in which the stimuli are identical within the entire region defined by the two locations of the square (see Figure 2), the standard square test measures the influence of image context outside that region.

**Figure 2.**
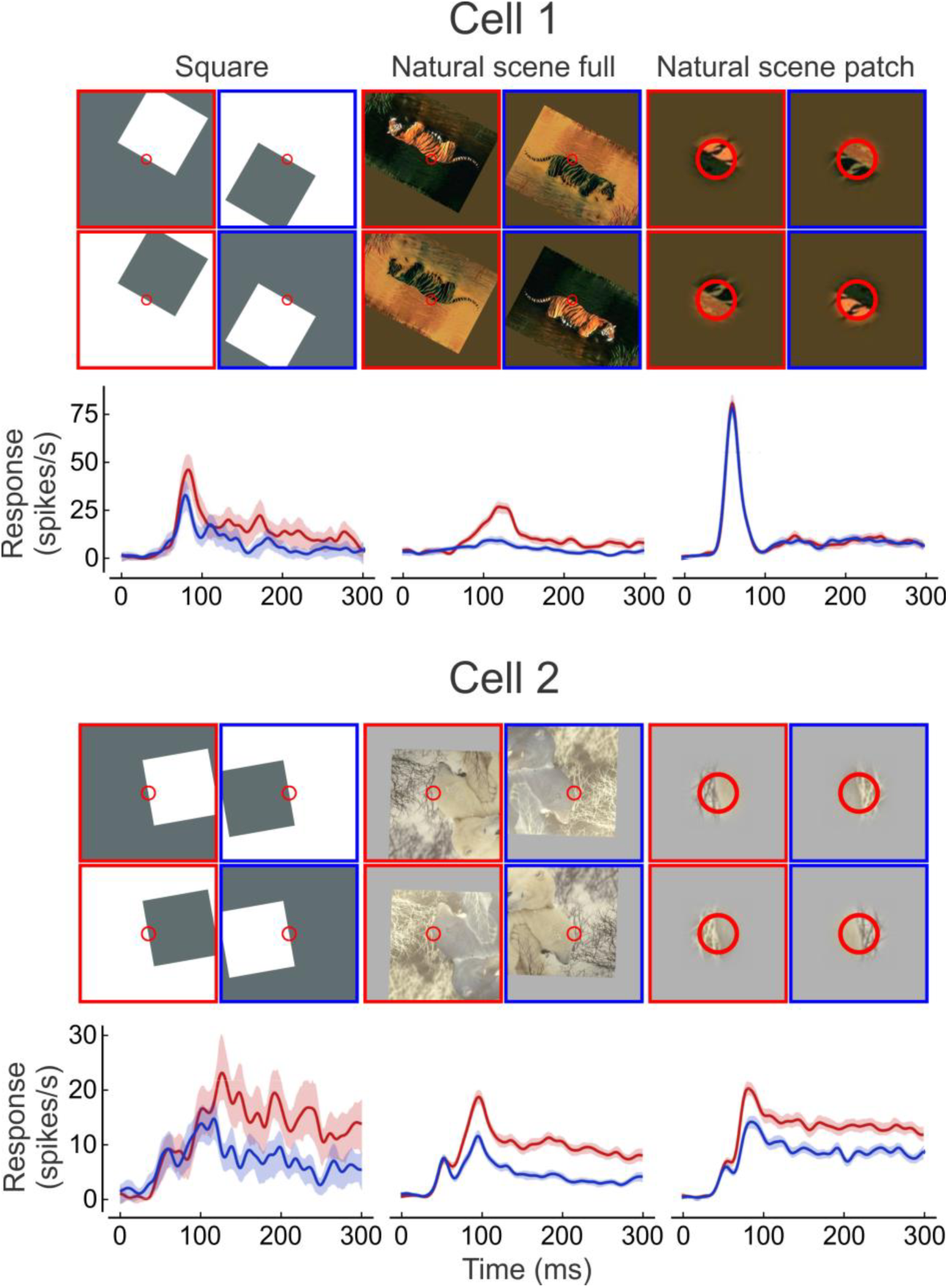
Responses from 2 example cells to square, full natural scene and patch of natural scene. Four frames are shown for each stimulus type with the two sides of border-ownership shown side by side, and the two contrast polarities at top and bottom. Red circles indicate location and approximate size of the CRF. The patch stimuli have been magnified for illustration purpose. Red frames indicate stimuli with objects on the preferred side and blue frames indicates stimuli on the opposite side. Only one of the presented scenes is shown in each case. Cell 1 was presented 44 scenes and cell 2 was presented 177 scenes. The mean temporal responses after the onset of the stimulus are plotted below with the corresponding colors, shading indicates 95% confidence intervals.

Our goal in this study was to test neurons in early visual cortex with occluding contours in natural scenes and see if they can signal which is the object side. How can we measure border-ownership signals elicited by natural scenes? In principle, we used the same strategy as in the standard square test. We took a sample of digitized photographs of natural scenes from a standard database (The Berkeley Segmentation Dataset, Martin et al., 2001). As humans, we understand the images and can thus identify objects and their contours. We placed an occluding contour across the CRF center of a neuron, rotated the image so that the contour matched the neuron’s preferred orientation and recorded the response for this orientation, and again after rotating the image 180°, keeping the contour in the center of the CRF. The rotation brings the object to the other side, thus reversing the direction of border-ownership in the receptive field. However, with natural contours the situation is more complicated than with the edge of a square, because these contours are generally not straight, and the regions on both sides of the contour are not uniform. Therefore, it is impossible to compare two situations that are locally identical, as we do in the standard square test. If the contour in the CRF is curved one way in the first presentation, it will curve the other way after the rotation. Even for a straight contour, we cannot make the two tests locally identical, because the regions adjacent to the contour are not uniform. A region may have texture or contain a luminance/color gradient. Curvature, texture and gradients are effective in determining perceived border-ownership in human vision (e.g., the concave side of a curvature is more likely to be the object side (Kanizsa, 1979), image structures that terminate at the contour are likely to be background (von der Heydt et al., 1984; Heitger et al., 1992; Heitger et al., 1998), and if one side shows a gradient perpendicular to the contour it is likely to be the object side (“extremal edge”, Palmer and Ghose, 2008). Because we cannot control these factors completely, we can only estimate border-ownership selectivity by testing each neuron with many different images. To get a better handle on the influence of local features, we expanded the factorial design by adding two dimensions of stimulus variation, “patch” and “edge-contrast-polarity”.

First, in addition to presenting the entire image we also presented a patch covering the CRF in isolation so that we could separate the effects of local and global factors. The stimulus presented within the CRF was identical in both the entire image and patch. Outside of the CRF, the patch blended into a uniform colored surround. The surround color was set to the mean of the color values within the patch. By rotating the patch 180° we determined the influence of the local border-ownership cues in isolation.

Second, to control for the effect of edge contrast polarity we devised a color inversion scheme analogous to that of the standard square test. This is important because contrast polarity is sensed by the CRF, which drives the neuron, and can strongly affect the strength of responses; in V2 and the supragranular layers of V1 about 50% of neurons are contrast polarity selective (Friedman et al., 2003). To control for this factor, we created color inverted images that essentially reversed the local edge contrast within the CRF. The test contour divides the patch into two roughly equal regions, and we calculated a transformation in 3D color space that flipped the mean colors of the two halves of the patch. The color values across the entire image were transformed this way. This transformation is analogous to flipping figure and ground colors in the standard square test, except that it is based on the mean colors in the vicinity of the CRF. One could of course perform the border-ownership test with only the original color images and determine the main effect of border-ownership. The contrast polarity would then be a random factor. Using color-inverted images allowed us to include contrast polarity as a covariate which greatly improved the power of the analysis.

Each neuron was tested with many different natural scenes (10-177, mean 43) and with the 2 sizes of squares in the standard square test. In early experiments we selected pieces of contours that matched the preferred orientation of the neuron under study approximately (±5°) and rotated the image by a small angle to match preferred orientation exactly. Thus, we kept the images upright except for a small rotation. With this method we had to use a large number of images, each with multiple test points. At later stages, we rather used a smaller subset of images/scene-points which enabled us to test the same set of scene points in every neuron. This implied larger rotations (±90°) to match the neurons’ preferred orientations. This selection of test points also excluded high local curvature and contour junctions, while the earlier set of images included such complex situations. Analysis of the data showed no difference between these two experimental schemes. Specifically, there was no significant difference between the firing rates in response to upright image and inverted images (p=0.87, two-sided Wilcoxon signed-rank test). We therefore pooled the results obtained with the two schemes.

We obtained complete sets of data from 140 V2 neurons of three monkeys, which we will refer to as FR (53 neurons), BE (50 neurons), and GR (37 neurons). Of these, 88 cells (63%) had a significant (p=0.01) effect of border-ownership in the standard square test, and 65 (46%) had a significant effect of border-ownership for objects in the full images of natural scenes. The border-ownership effect from the global context, as measured by the interaction of Patch with Border-ownership, was significant in 38 cells (28%). Because testing natural images was sometimes skipped if border-ownership selectivity was not obvious in the standard test, these percentages may overestimate the true frequency of border-ownership selectivity. E.g. our sample has 63% of the cells selective with the standard square test, whereas previous studies found about 50% (in V2).

As mentioned above, the 180° image rotation did not affect the mean strength of responses across the population. Thus, the border-ownership effects in natural scenes are not the result of a response difference between upright and inverted scenes.

### Contours in natural scenes versus edges of squares

How do the border-ownership signals for complex natural scenes compare with those for the much simpler standard square test displays where the object is a figure surrounded by uniform color? Because the standard square test measures the influence of context, it was of interest to compare its results with the context effect in natural scenes. In 33 cells, both the standard square test and the context of natural scenes produced significant border-ownership modulation (p=0.01). Of these, only 2 cells had opposite preferences. That is, 94% of the cells were found to signal side-of-object consistently in the two tests.

The post-stimulus time histograms of the responses from two example cells are illustrated in Figure 2, along with an example scene for each (in total, cell 1 was presented 44 scenes, cell 2, 177 scenes; the histograms for natural scenes are means across scenes). The conditions in which the square or natural object was on the cell’s preferred side are demarked with red and the conditions in which they were presented on the cell’s non-preferred side are demarked with blue. The preferred side is defined consistently for each cell (using the square test as the standard). Cell 1 (Figure 2A) produced a significant (p=0.01) mean border-ownership signal with the full-image displays, which can be seen in the higher mean firing rate when the objects appeared on the preferred side (red) compared to the non-preferred side (blue), but not with the patches. Cell 2 (Figure 2B) produced a significant mean border-ownership signal in both full-image and patch conditions. The significant signal for patch indicates that the cell was sensitive to local figure-ground cues (e.g., curvature, edge luminance profile, texture differences etc.). As we shall see, the behavior of Cell 1 is more representative of the population results; while there were 35 cells with significant border-ownership signals in the patch condition (p=0.01), only 20 of these cells (57%) were consistent between the patch condition and the standard square test (not significantly greater than chance, p=0.05, one-tailed binomial test).

To show how the border-ownership signals for natural scenes and square relate across the population of cells, we plotted their mean signal across natural scenes versus their signal for the standard square test (Figure 3). For each neuron, the border-ownership effects in both tests were determined by linear models and divided by the square-root of the residual variance from the model of the square test (see Methods). Because the model of the square test includes all variables and interactions, its residual variance reflects only random response variation. Thus, the figure represents the border-ownership signals of each neuron in multiples of the standard deviation of the random variation of its responses. Note that all signals of each neuron were scaled by the same factor to allow comparison between conditions.

**Figure 3.**
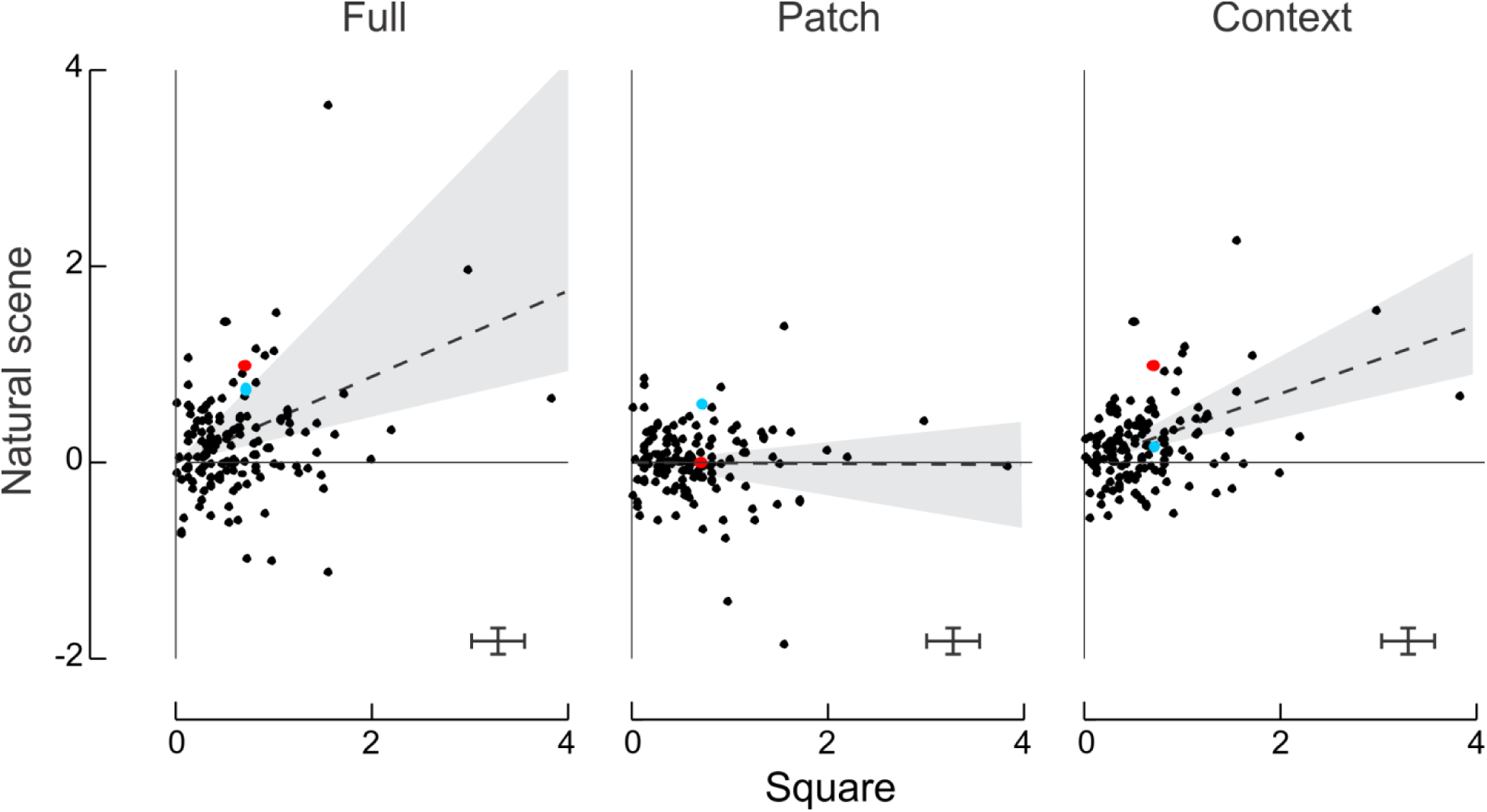
Comparison of the border-ownership signals produced by natural scenes and squares. For each of the 140 V2 cells studied, the border-ownership effects of full image and patch, and the context influence, are plotted against the border-ownership effect of squares. The effects were measured by linear regression performed on square-root transformed spike counts (see Methods). Error bars bracket the range of 6 times the mean standard errors of estimates. The slopes of the lines, determined by minimizing the orthogonal squared deviations, indicate the relative strengths of border-ownership signals for the natural scene stimuli relative to squares across the population. Shaded areas and dashed lines indicate 95% confidence intervals. Colors mark the example neurons of Figure 2 (red, Cell 1, blue, Cell 2).

To compare the strength of signals for contours in natural scenes with that for edges of squares, we calculated the regression lines through the origin minimizing the orthogonal squared deviations (see Methods). Minimizing absolute deviations instead of squared deviations produced similar results, as did fitting the direct effects rather than normalized effects. The positive slopes of the regression lines indicate that there was overall agreement between the results for the full images of natural scenes and the standard square test, and between the context effects of natural scenes and the standard square test. The values of the slopes also indicate that the border-ownership signals for natural scenes tended to be weaker than those for the squares. For the full natural scenes, the relative strength was 44% and for the context influence alone it was 35%. The slope for the patches was not significantly different from zero, indicating that the effects of local border-ownership cues in isolation were not consistent with the border-ownership signals evoked by the squares. Note also that the context signals were less scattered than the full-image signals. Apparently, separating the influences of local cues from the context effects reduced the variance, presumably because the effects of local cues did not correlate with the border-ownership signals for edges of squares.

Computing the slopes for the sub-population of cells that were border-ownership selective (P<0.01) for both, natural scenes and squares, the strength of the border-ownership signal for natural scenes was 72% of that for squares (95% CI: 36 to 148%), the influence of the context was 59% (CI: 33 to 89%), while the local cues still had no effect (2%, CI: -12 to 27%).

The scatter in the plots of Figure 3 shows that the relative strength of the border ownership effects for natural scenes and square figures varied greatly between neurons. This is the reason for the wide confidence intervals of the fitted slopes. The scatter in these plots is not simply the result of random response variation. In fact, the confidence intervals of the effects in the individual neurons were quite small: error bars in the Figure indicate 6 times the mean standard errors (^~^99% CI). Rather, the scatter indicates genuine differences between neurons in processing the two kinds of stimuli. Zhou et al. (2000) noted this variation when comparing border ownership selectivity between different configurations of geometrical shapes and attributed it to differences between neurons in the way they evaluate the available cues. Our results discussed in the next section confirm this conjecture.

### Variation of border-ownership signals across scenes

Above we examined the strength of border-ownership signals looking at means across scenes. Perhaps the most intriguing question is how consistent the signals are across the different scenes. In Figure 4 we have plotted, for the two example cells, the border-ownership effects of the individual scenes, sorted by size, for the full-image and patch conditions and for the context influence (black curves). The intersections of the curves with the abscissa are shifted to the right of the center because the majority of scenes produced positive effects, except for the patch condition in cell 1 (Figure 4A) which produced no consistent effects.

**Figure 4.**
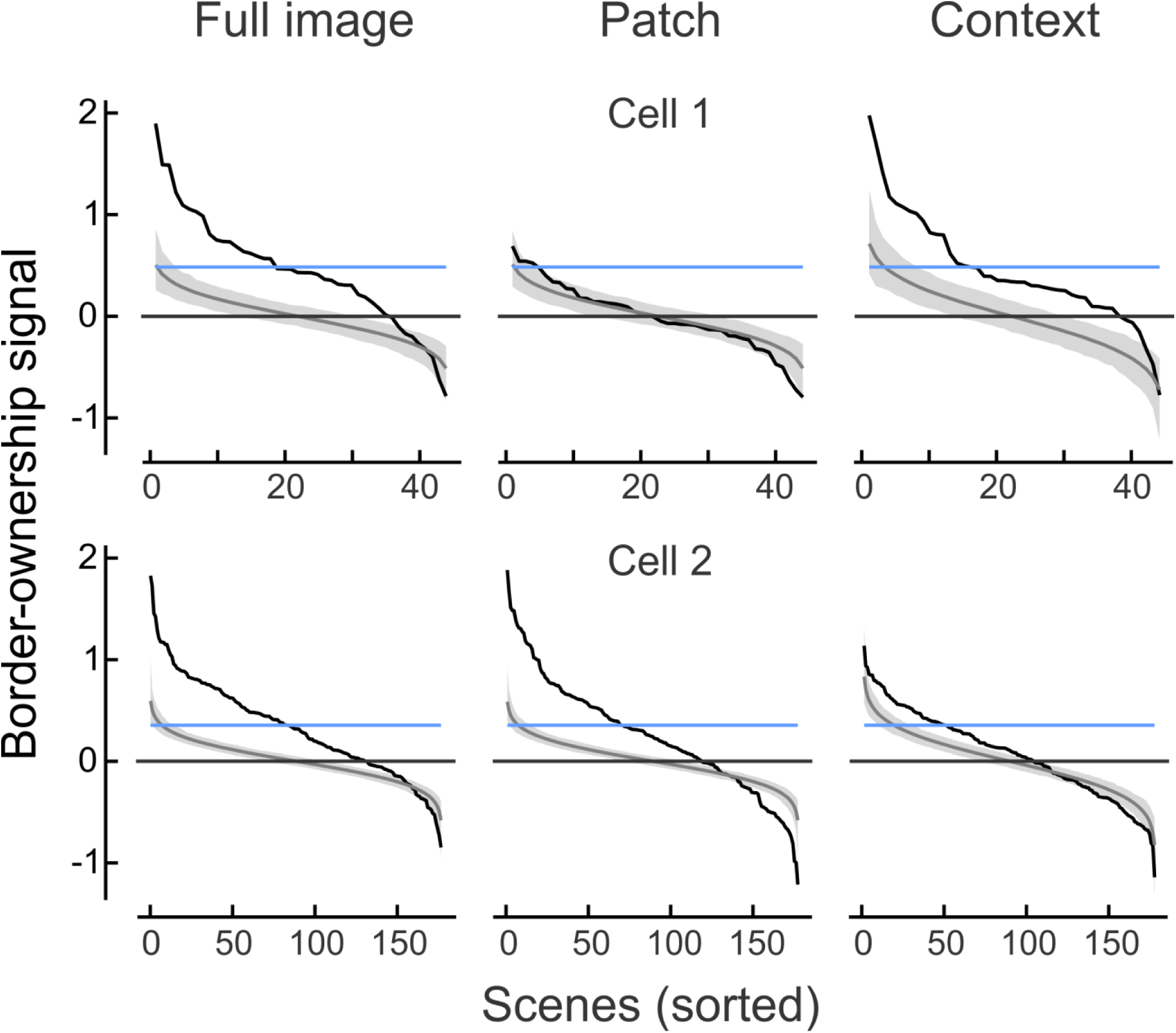
Variation of border-ownership signals across scenes. Data from the same cells as in Figure 2. Scenes were sorted, in decreasing order, by strength of border-ownership signal. Black lines represent signals for full image, patch, and context. Gray bands show the 95% confidence intervals of the sorted effects obtained under the null hypothesis (no border ownership selectivity). Horizontal blue line indicates the border-ownership signal for squares.

These plots indicate considerable variation of the effects across scenes, with strong positive effects (consistent with the standard square test) for some scenes and weak or even negative effects for others. Of course, some of this variation is caused by the random variation of responses. Determining the effects for the individual images includes substantial amounts of random variance because the number of responses to each scene was small. We determined the effect of random variation by generating for each recorded neuron a large number of surrogate neurons with the same random variance, but no border-ownership selectivity, and sorting the predicted ‘border-ownership effects’ of each surrogate neuron (see Methods). The mean across the surrogate neurons represents the null hypothesis that border-ownership has no influence (gray trace in Figure 4, shading represents 95% confidence limits). The null hypothesis curve of course intersects the zero line near the center as about half the values must be positive and half negative. The slope of this curve is entirely the result of sorting random variations.

Comparison with the null-hypothesis curve suggests that some of the negative values of the data are the result of random response variation. Particularly in the case of the patch condition in cell 1 (Figure 4A), the null hypothesis might explain nearly all the variation of border-ownership effects between scenes, indicating that the pictorial cues within the CRF had little, if any influence. Thus, in this neuron, the border-ownership signals in the full-image condition were driven mainly by the image context.

The random variation of responses affects the apparent consistency of the border-ownership effects across scenes. Because these variations are not correlated with the border-ownership effects, they do not systematically affect the estimate of its mean, but they tend to lower the proportion of consistent values: in the limit, when the random variance is large compared to the mean, the proportion of positive values approaches 0.5 (chance). More specifically, the measured distribution of border-ownership effects is the convolution of the true distribution of border-ownership effects with the noise distribution. The latter equals the distribution of simulated border-ownership effects under the null hypothesis. Thus, in principle, we can recover the true distribution by deconvolution (see Methods). Validation on simulated neurons with known border-ownership consistencies showed that the deconvolution provided a better estimate of consistency than using the raw border-ownership effects while still tending to underestimate the true consistency.

Figure 5 shows the distribution across neurons of the proportion-consistent for the full-image condition as derived from the raw data (top) and as corrected by deconvolution (bottom). The corrected results show that 13 cells were over 80% consistent across scenes, and 3 were over 90% consistent. Example cell 1 (Figure 2) was 83% consistent (p=6·10^−4^, testing proportion against 0.5, Bonferroni corrected for testing the 65 proportions) and cell 2 was 79% consistent (p=6·10^−14^, ditto). When an occluding contour is placed in the receptive field of such a neuron it signals the object side correctly with fairly high probability.

**Figure 5.**
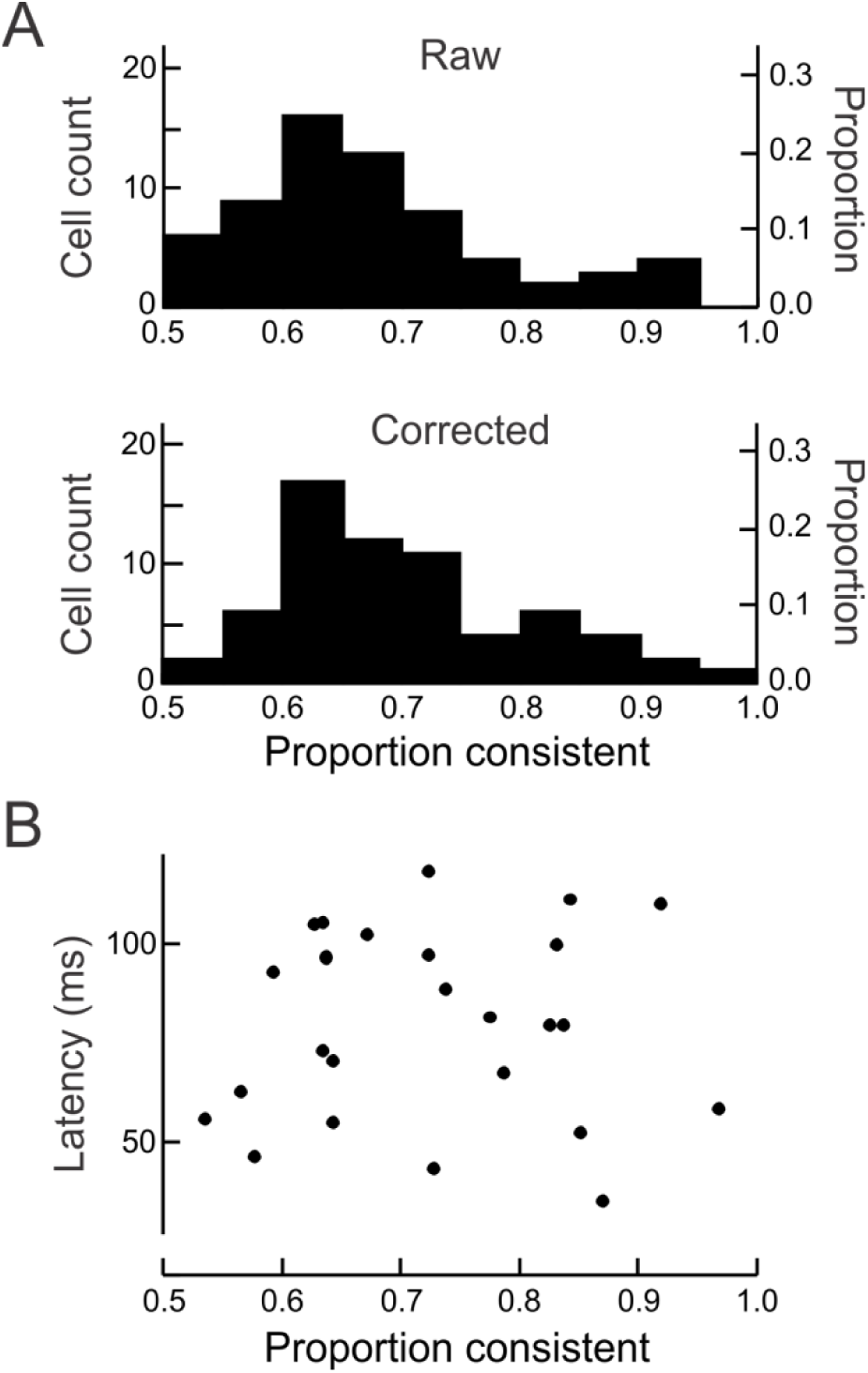
Consistency and latency of neurons in signaling border-ownership for natural scenes. **A**, ‘Raw’ shows the distribution of the proportion of scenes for which a cell gave the same sign of border-ownership signals. ‘Corrected’ shows the distribution of the proportion after correction for random variation (see Results for details). Cells are selected for significant (p<0.01) effect of border-ownership in full natural scenes (N=65). **B**, Estimates of the latencies of the border-ownership signals for natural scenes in the individual cells plotted as a function of their consistency. The latency of the border-ownership signals did not increase with consistency.

### Time course

How can neurons at this low level in the visual cortex signal consistent figure-ground interpretations for natural scenes? Solving the figure-ground problem in complex images clearly requires some understanding of the image. Such understanding might be based on object recognition. Is it possible that the border-ownership signals for natural scenes are the result of feedback from shape selective neurons in IT cortex? Because IT neurons only start responding about 80ms after stimulus onset and become shape selective even later (around 150ms, Brincat and Connor, 2004), border-ownership modulation for contours in natural scenes would then emerge with a corresponding delay, and much later than the border-ownership signals for edges of squares which appear around 68ms after stimulus onset (Zhou et al., 2000; Sugihara et al., 2011). Surprisingly, this was not the case.

We computed the average post-stimulus time histograms for the different stimulus conditions using the data of the 31 cells that showed significant (p<0.01) effects of border-ownership for both squares and natural scenes, excluding the 2 cells in which the effects were opposite (because the BO signal is the difference between preferred and non-preferred side responses, cells with opposite preferences cannot be included because choosing one or the other side would bias the result towards either natural scenes or squares). A comparison of the histograms (Figure 6A) shows that the border-ownership signals emerged virtually at the same time for natural scenes (red line) as for squares (black dashed line).

**Figure 6.**
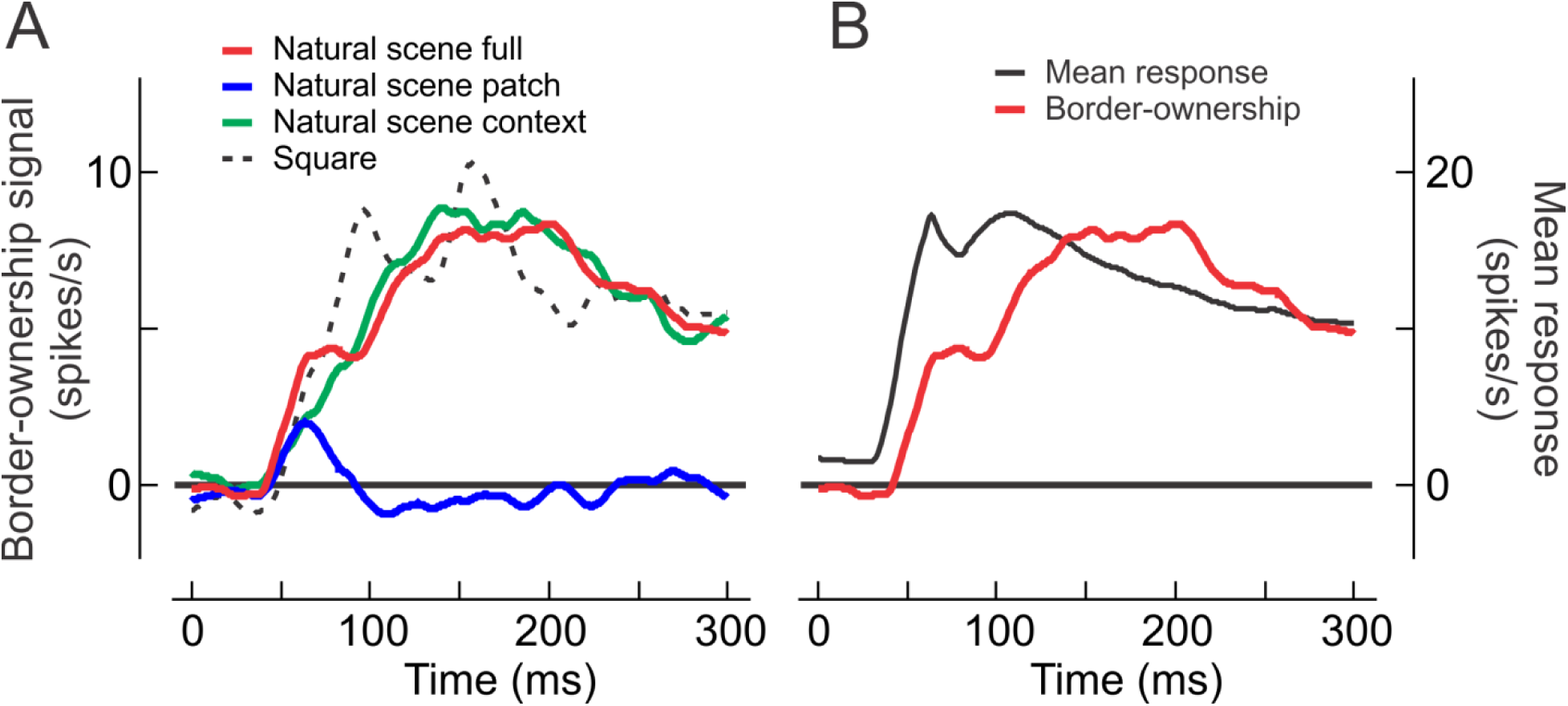
The time course of border-ownership signals for squares and objects in natural scenes. **A**, The curves show smoothed post-stimulus time histograms of the mean border-ownership signals, averaged over cells, for full natural scene (red), natural scene patch (blue), natural scene context (green) and square (black dashed). The border-ownership signal for natural scenes and its context component emerge at the same time as the signal for square figures, but rise slightly more slowly. The patches of natural scenes produce only a small transient signal. **B**, Comparison of the time course of border-ownership signal (red) and mean response (black) for contours of natural scenes. Note the short interval between response onset and rise of the border-ownership signal.

We also plotted the border-ownership modulation produced by the image patches (blue) and the context effect, i.e., the difference between full-image and patch effects (green). The image patches produced only a small transient positive signal between about 40-90 ms which then disappeared, whereas the image context effect rose about linearly, peaking at 140 ms. The image context effect should be compared with the border-ownership signal produced by the square (here shown for the 8° size) which also depends entirely on the image context (the displays produced by flipping the square about the test edge and reversing the contrast are identical within a 16°×8° region centered on the CRF). Although both emerge at the same time, one can see that the curve for natural scenes (red) rises slightly more slowly than the curve for the square (black dashed). All border-ownership signals except those for the patch lasted until the end of the observation period (300 ms), decaying slightly after the peak.

In Figure 6B we compare the border-ownership signal for natural scenes (red, replotted from A) with the mean of the responses to preferred and non-preferred sides (black, right-hand scale). Note the short delay between response onset and border-ownership signal.

The results of Figure 6 are consistent with the scatter plots of Figure 4 in showing that the border-ownership signals, averaged over the analysis period, did not correlate between the patch and square conditions, and that the context effects of natural scenes correlate more tightly with the square signals than the full-image signals did. Apparently, the CRF produces consistent border-ownership signals only briefly after stimulus onset, but does not have a consistent effect later.

Table 1 summarizes the latencies for responses and border-ownership signals in the different conditions as determined by two-phase regression on the cumulative spike count histograms (Figure 7, see Methods). These latencies correspond approximately to the time point at half-maximal strength of the signal. The table shows a latency of 60 ms for the full image border-ownership signal and 73 ms for the context influence, compared to 44 ms for the response onset. The estimate for the response onset agrees with previous estimates for edge responses in V2 (Zhang and von der Heydt, 2010). Thus, assuming that the cortex computes the border-ownership signal from the V2 responses, this leaves only about 16 ms for computing the border-ownership signal for the full image, and 29 ms for evaluating the image context.

**Figure 7.**
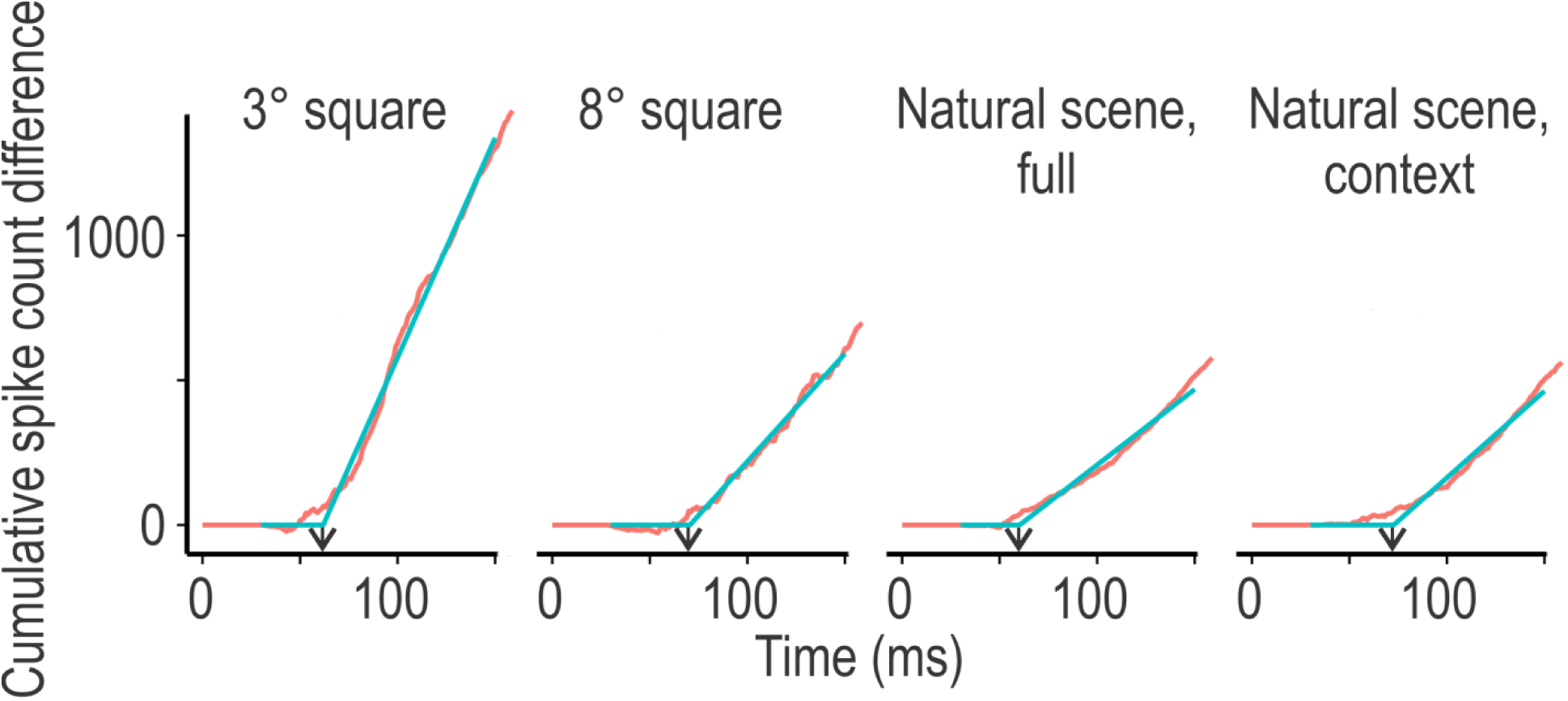
Determination of the latencies of the population border ownership signals. Red curves show the cumulative spike difference counts for small and large squares, the full natural scenes and the context component. The latencies were determined by 2-phase regression fits (blue lines, see Methods).

**Table 1.** Summary of the latencies of responses and border ownership signals. Latencies for onset response and border-ownership signal and their standard deviations for 33 border-ownership selective cells.

| Stimulus | Onset response (ms) | Border-ownership signal (ms) |
| --- | --- | --- |
| Square (3°) | $46 \pm 0.3$ | $62 \pm 3$ |
| Square (8°) | $46 \pm 0.3$ | $71 \pm 6$ |
| Natural scenes, Full | $44 \pm 0.1$ | $60 \pm 2$ |
| Natural scenes, Patch | $47 \pm 0.1$ | |
| Natural scenes, Context | | $73 \pm 4$ |

The short latency of the mean border-ownership signal for natural scenes is surprising considering the high consistency of the border-ownership effects across scenes in some neurons (Figure 5). Could the early part of the signal be contributed by less consistent neurons that produce strong signals, but only for a few scenes, whereas the later part of the signal comes from highly consistent neurons with longer latency (which would be compatible with feedback from IT)? Our data contradict this explanation: Cell 1 in Figure 2 was highly consistent (83%, 44 scenes tested) and produced border-ownership modulation with short latency (85ms). Across the population, there was no evidence of latencies being longer for highly consistent neurons (Figure 5B). There was no correlation between Fisher-transformed consistency index and latency (Pearson r=−0.04, p=0.85).

Taken together, our analysis revealed no indication of a gradual emergence of border-ownership signals for natural scenes that would correlate with the progression of shape selective responses in the ventral stream. The border-ownership signals pop up extremely fast, within less than 30ms after the onset of responses in visual cortex.

## Discussion

This is the first neurophysiological study of border-ownership selectivity of cortical neurons in natural scenes. Our main findings are that a subset of V2 cells consistently signal border-ownership in natural scenes, that the border-ownership coding for natural scenes is consistent with that for simple figure displays like a square in a uniform field, and that the border-ownership signals for natural scenes emerge at the same time as the border-ownership signals for simple figures, reaching their maximal strength only slightly later. We believe our results portray a picture of the neural activity that corresponds closely to how real scenes are represented in the visual cortex under natural viewing: Recording from neurons in awake monkeys during periods of fixation, we studied the responses of orientation selective neurons (which constitute over 75% of the neurons in V2, Friedman et al., 2003) by aligning their receptive fields with occluding contours in images of natural scenes. As a consequence, the neurons generally responded strongly to at least some of the images. Thus, we are looking at the pattern of activity across the population of neurons that would be strongly activated by objects in natural scenes. These neurons presumably provide the information that subsequent object processing centers rely on.

### Robust mechanisms

The border-ownership signals showed a remarkable degree of consistency both across the population of neurons and across scenes within the individual neuron. Among the neurons that were significantly modulated by border-ownership for both, natural scene contours and borders of squares (about ¼ of our sample), the mean border-ownership signals in the two conditions were consistent in 94% of the neurons. Comparing the responses of individual neurons across large samples of different natural scenes, we found that some neurons were 90% consistent across scenes. These V2 neurons perform much better than the simulated neurons in a somewhat simplistic model (Sakai et al., 2012) which reached only 67% consistency on images of the Berkeley Segmentation Dataset. We are not aware of other neural models that have been evaluated on images of natural scenes. The consistency across scenes rules out the possibility that the mean effects of border-ownership were the result of occasional conspicuous features or configurations. Finding such a high degree of consistency in neurons of a low-level visual area like V2 is surprising.

Our results of course apply to our specific selection of scenes and test points which is somewhat arbitrary. However, as the example set in Figure 1 shows, our test contours included a large variety of situations: borders between regions of different contrasts, colors and textures, and borders with luminance/color gradients. Also the shapes and sizes of the objects varied greatly. Thus, the observed consistency of border-ownership coding shows that border-ownership coding reflects mechanisms for detection of “objectness” that are highly sophisticated and robust.

**Figure 1.**
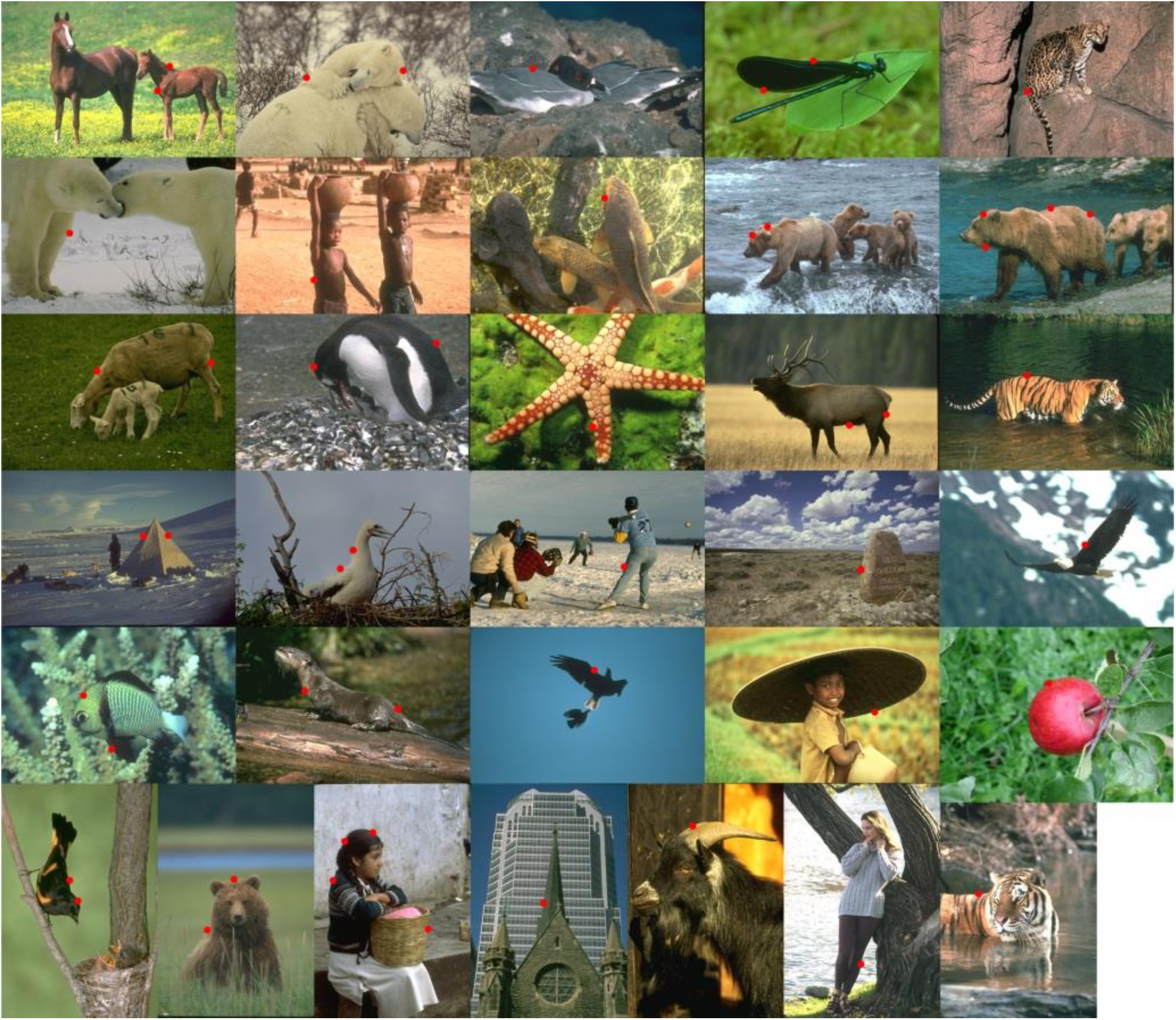
Example scenes used for testing border-ownership selectivity of neurons. Selected points on occluding contours (marked here by red dots for illustration) were centered in the receptive field of the neuron to be studied. The examples show the most frequently used scenes, with the number of neurons tested per scene ranging from 90 (top left) to 10 (bottom right). All of the images used were from the Berkeley Segmentation Dataset, except for the image of the apple on the lower right.

There are many local cues that can influence our perception of border-ownership (Peterson and Salvagio, 2010). Our results from the image patches show that the CRFs did not contribute much to the border-ownership signals, except for the brief period from about 50 to 90 ms after stimulus onset (Figure 6). The weakness of border-ownership signals from patches is consistent with perceptual and computational studies (Fowlkes et al., 2007) showing that human performance strongly depends on the size of the image patch and would be poor or at chance level for the small patch sizes we used, especially since we excluded regions of high curvature and contour junctions. It is important to note that finding weak effects in the patch condition does not mean that local cues have little influence on the border-ownership computation. The patch generates a brief signal which is later discarded as there is no confirmation from the context. Most likely, the border-ownership mechanisms integrate local cues all along the contours as proposed in the grouping cell model discussed below.

### Fast processing

Perhaps the most surprising result is the early onset of border-ownership signals in the natural scenes test. Using two-phase regression on the cumulative differential spike count histograms we estimated signal latencies (corresponding approximately to the time point when the border-ownership signal reaches half-maximal strength) of 60 ms for contours in natural scenes as a whole, and 73 ms for the component contributed by the image context (Table 1). These latencies are essentially the same as the latencies of the border-ownership signals for edges of squares (62 and 71 ms for sizes of 3° and 8°, respectively). Because neurons in area V2 only start responding around 45 ms (in all conditions) this leaves only 16 ms processing time in the case of full images and 29 ms for the context influence.

### Searching for an explanation

Explaining the robust and fast figure-ground signals is a challenge for the theory and modeling of visual cortical function. The spread of feed-forward connections, which account for the CRF, is too small to account for the context integration (Angelucci et al., 2002; Zhou et al., 2000). Also lateral propagation of signals within the cortex (V1 or V2) is unlikely to explain these results, because the conduction velocity of intracortical (“horizontal”) fibers is too slow to transmit the necessary context information across these large retinotopic representations, an argument that is based on an analysis of border-ownership signals for edges of squares, where the cortical distances can be determined exactly (Craft et al., 2007; Sugihara et al., 2011). By extension, given the similarity of the time course between the border-ownership responses to natural scenes and squares (Figure 6), it seems unlikely that lateral propagation would underlie the natural scene data. This seems to rule out models based on intracortical propagation of signals (Zhaoping, 2005).

The short latency of the border-ownership signals for natural scenes also seems to rule out mechanisms tuned to specific object shapes in inferotemporal cortex as an explanation, because neurons there only start responding around 80ms after stimulus onset and become shape selective only around 150ms (Brincat and Connor, 2004). Also applying a classifier-based readout technique to the responses of many IT neurons yields valid object categorization only after ^~^100ms (Hung et al., 2005). Thus, any top-down influence from specific object recognition mechanisms would arrive much later than the observed border-ownership signals.

By elimination, we conclude that border-ownership coding must involve back-projections to V1 and V2 from hypothetical higher-level mechanisms that are fast enough to generate consistent signals within a few tens of milliseconds. Large context integration occurs in a number of cortical areas that are only a few centimeters away, and, because the forward- and back-projection loops consist of white matter fibers which have high conduction velocity, this hypothesis can explain the critical observation of wide context integration with short latency.

This is the basic idea of the “grouping cell model” proposed earlier (Craft et al., 2007; Qiu et al., 2007). This model explains border-ownership coding for the relatively simple displays of geometrical figures, such as a geometrical figure surrounded by a uniform region (Zhou et al., 2000), pairs of overlapping figures (Zhou et al., 2000; Qiu et al., 2007), and the concave part of C-shaped figures (Zhou et al., 2000). In this model, neural responses from edge selective neurons (simple or complex cells in areas V1 and V2) feed into grouping cells (“G-cells”) at a higher level, which, by feedback, set the gain of the same edge neurons, thus modulating their visual responses. The feedback modulation makes these edge neurons border-ownership selective (“B-cells”). The G-cells receive additional bottom-up input from end-stopped cells responding to T-junctions which suggest occlusion and indicate the direction of occlusion (Heitger et al., 1992). Thus, G-cells accumulate evidence from local cues distributed along the contour. However, an isolated patch containing such a cue would not be sufficient to keep a G-cell active, which explains the brief duration of the neural signal in the patch condition (Figure 6). Analyzing spike correlations between simultaneously recorded neurons Martin & von der Heydt (2015) found increased synchrony precisely between those neurons that, according to the model, receive common input from grouping cells, in strong support of this model.

In this model (Craft et al., 2007) the grouping is based on very simple fixed summation templates. G-cells sum “co-circular” edge signals. Per design, they respond best to objects of compact shape, and bias the border-ownership responses even when only a few roughly co-circular edges are present, in agreement with the neurophysiology (Zhang and von der Heydt, 2010). Whether this simple model would also explain the present results on natural scenes needs to be seen.

### Border-ownership and saccadic eye movements

Understanding the object structure of scenes from images is a fundamental task of vision. Our results show that this task is performed to some extent by extremely fast mechanisms that do not depend on feedback from object-recognition centers. The same mechanisms might underlie the ultra-rapid object detection found by Thorpe et al. (Kirchner and Thorpe, 2006; Crouzet et al., 2010). When two scenes are simultaneously flashed in the left and right hemifields, human observers can reliably make saccades to the side containing an animal in as little as 120 ms. Saccades to faces are even faster. As suggested by Martin and von der Heydt (2015) the activation of grouping circuits corresponds to the formation of “object files” or “proto-objects” postulated in perceptual theories (Kahneman et al., 1992; Rensink, 2000). The grouping circuits are thought to provide the structure for object-based attention (Qiu et al., 2007; Craft et al., 2007; Mihalas et al., 2011) which is closely related to saccade planning. Thus, the fast border ownership signals for natural scenes demonstrated in the present study might reflect the formation of proto-objects which also enable the system to make fast saccades to objects.

## Materials and Methods

We studied neurons in the visual cortices of three male rhesus macaques (*Macaca mulatta*). All procedures conformed to National Institutes of Health and United States Department of Agriculture guidelines as verified by the Animal Care and Use Committee of Johns Hopkins University.

### Preparation

Three small head posts for head fixation were implanted in the skull and a recording chamber was placed over the visual cortex of each hemisphere under general anesthesia.

### Recording procedures

Isolated neuronal activity was recorded extracellularly with glass-coated platinum-iridium microelectrodes (PtIr 0.1 mm diameter, etched taper ^~^0.1, impedance 3-9MΩ at 1kHz) that were inserted through the dura mater. A spike time detection system (Alpha Omega MSD 3.22) was used.

Most of the V2 neurons were located in the lunate sulcus after passing through V1, the remaining neurons were located in the lip of the post-lunate gyrus. The eccentricities of the receptive fields ranged from 0.53 to 4.9 degrees of visual angle (median of 2.5).

### Stimuli and Experimental Design

#### Stimulus display

The stimuli were presented to the monkeys with either a 21-inch EIZO FlexScan T965 or a ViewSonic G220fb color monitor. Both had the refresh rate set to 100 Hz and resolution to 1600 × 1200. The monitors were viewed at a distance of 1 meter and subtended 21 × 16 degrees of the visual field. The luminous outputs of the monitors were linearized. The images of the natural scenes were gamma corrected with the equation 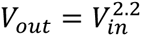 for the linearized displays.

#### Recording of gaze direction

The direction of gaze was recorded for one eye by corneal reflection or pupil tracking using an infrared video system (Iscan ETL-200) that was aligned with the axis of the eye via an infrared-reflecting mirror. The system recorded direction of gaze with a resolution of 0.08 × 0.16 degrees of visual angle, although the accuracy was lowered by noise and drifts in the signal.

#### Behavioral design

All data were collected using a fixation paradigm. Monkeys were given a juice reward for keeping the eye position signal within 1 visual degree of the center of a fixation point for 3-4 seconds.

#### Mapping procedures

After isolating a cell, its CRF was manually mapped with bars, drifting gratings, and/or rectangles, depending on the CRF properties of the neuron. Color and orientation were varied to determine the optimal stimulus. The manual mapping was typically confirmed by a position test that systematically moved a luminance step-edge or bar in randomized order.

The orientation preference was first tested in steps of 30 degrees (from 0 to 330 degrees). Often the orientation was fine-tuned using smaller steps over a more focused range. Only cells that were orientation selective were included in the results. This was determined by using the orientation modulation index when orientation tests were presented or by manual mapping when the orientation preference was obvious. The orientation modulation index was calculated by:

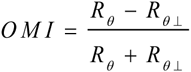

where *R*_*θ*_ and *R*_*θ*⊥_ are mean responses to the preferred and orthogonal orientations. Only cells that had an orientation modulation index of at least 0.20 were included in the analysis.

#### Standard square border-ownership test

The border-ownership signal was first assessed using the standard square border-ownership test as described previously (Qiu and von der Heydt, 2005). This test presents a square with one of the edges centered over the CRF of the cell and the square rotated such that the orientation of the border matches the cell’s preferred orientation. The square and the background are defined by two colors: gray (28 cd/m^2^) and the preferred color of the cell (or a color which elicited a strong response). The assignment of each of these colors to the square and the background were counterbalanced, in order to separate out the effect of local contrast from the border-ownership coding. Two sizes of squares were used: three and eight degrees. A uniform background with the mean of these two colors was displayed between stimulus presentations.

For the standard square border-ownership test and the natural scene border-ownership test (described later) multiple stimuli were presented within a behavioral fixation period. Each stimulus was presented for 300 ms and a uniform screen was shown for 200 ms between the stimulus presentations whose color was set to the mean stimulus color within the CRF.

#### Natural scene border-ownership test

We used images from the Berkeley Segmentation Dataset 300 (Martin et al., 2001) and one additional image^1^. Before data collection, we labeled many points in the images that lay on the occluding contours of objects, recording position, orientation of contour and side of the occluding object. Each neuron was tested with a number of images aligning one of these points in the center of the CRF. Custom software was used to assist in accurately positioning the points. We call each test point in an image a “scene”. Not all of the recorded neurons were presented with the same scenes. Figure 1 shows 51 points denoted in the 32 images that were used most frequently in the experiments. The exact subset varied between monkeys, because some scenes seemed to distract a given monkey more than others from the fixation task leading to frequent abortion of trials. Typically only one point per image was tested, although sometimes there were more. In early experiments, we selected, for each neuron, points where the contour orientation was within 5 degrees of the neuron’s preferred orientation and rotated the images so as to match the preferred orientation exactly. In later experiments, we selected a subset of scenes (shown in Figure 1) and applied arbitrary rotation. This enabled us to test the same scenes across multiple cells.

In the initial experiments, when we selected points based on how closely their contours matched the recorded neuron’s preferred orientation, we included more complex images. However, when we later concentrated on a smaller subset of points (shown in Figure 1), we avoided points with high curvature, contour junctions, and thin objects, where both sides of the objects could fall within the receptive field.

For each scene, we manipulated the side-of-object, the contrast polarity, and the image context (Figure 2),

*Side-of-object* – In order to manipulate the position of the occluding object (relative to the CRF, either preferred or non-preferred side) for a given scene, the image was rotated 180 degrees about the center of the CRF.

*Contrast polarity* – Rotation of the images by 180 degrees changes the local contrast polarity within the CRF, since the local contrast rotates with the image. In order to control for this, we performed a transformation of the RGB color-space that inverted the mean colors within the CRF on either side of the border. The color inversion transformation we used for a given scene (scene point), *s*, is:

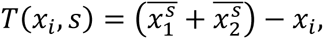

where 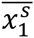 and 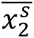 are the average colors on each side of the border within the CRF and *x_i_* is the color of the pixel that is being transformed. This is similar to the common color inversion transformation, however, the color-space is inverted around the mean colors within the CRF instead of the mid-point of the color-space.

*Context* – In order to separate the effect of the local stimulus within the CRF (which is driving the response) from the modulation by the context, we showed both the full images and local patches that covered the CRF. The stimulus within the approximate receptive field radius, **r**, was exactly the same, while outside this radius, the stimulus faded into the background using the complementary error function:

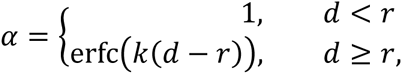

where *d* is the distance from the center of the receptive field in pixels and *k* was set to 1.8.

Each of these factors (side-of-object, contrast polarity, context) is binary such that there are a total of 8 variations of each scene. All of the stimuli were presented in pseudorandom order.

The background color surrounding the full images and patches, and the display color during the interstimulus intervals for a given scene, was the average color within the area of the CRF,

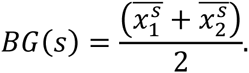

This definition of background color for each scene is analogous to the definition of the interstimulus color used in the standard square test.

### Data analysis

#### Analysis of spike counts

We used linear models to analyze each cell separately. The spike counts between 40 and 300 ms of stimulus onset were transformed by the Anscombe transform 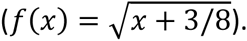. This transform approximately converts Poisson data to Gaussian distributions. Repeated measures ANOVA was used to measure the effect of border-ownership and its significance. A three-way fixed effects model was used for the standard square test data (border-ownership × edge-contrast-polarity × size), and a four-way model for the natural scenes data (border-ownership × edge-contrast-polarity × patch × scene-id). All of the factors were binomial except for scene-id which had multiple levels and could also vary between cells. Both analyses were based on factorial design and included all interactions. The context influence on the border-ownership signal in natural image was defined as the interaction between the border-ownership and patch.

To determine consistency of border-ownership signals and compare their relative strengths across different stimulus conditions we plotted the border-ownership effects of the full images, the patches, and the context alone, as a function of the border-ownership effects of the standard square test, and in each case fitted a line through the origin using orthogonal least-squares regression. The border-ownership effects were calculated using the linear models described above. Because firing rates and reliability of responses vary between cells, we plotted the effects in each cell were divided by the square-root of its error variance as obtained from the model fit to the standard square test data. Because this model contains all experimental variables and their interactions the error variance from it reflects the variation of responses between repeated presentations of the same visual conditions. This normalization is equivalent to weighting cells by their reliability. Orthogonal regression was used to treat both variables symmetrically. The fit was forced through the origin because the sign of the border-ownership signal for each neuron is arbitrary.^2^ This ambiguity means that the fit must be invariant against reflecting any data points about the origin. Thus, the fitted line must pass through the origin. (For the figures we reflected the data points at the origin so as to make the effects from standard square test positive, thus plotting all results in two quadrants.) The 95% confidence intervals for the slope of the line were calculated by bootstrap, resampling cells with replacement, using the R boot package (Davison, 1997; Canty and Ripley, 2015).

#### Variation of border-ownership signals across scenes

To illustrate the variation of border-ownership signals in a neuron across scenes we sorted the scenes by the strength the border-ownership effect, separately for full images, patches, and context. We then generated the distribution of border-ownership effects that would result if border-ownership had no influence (null hypothesis). We generated the null hypothesis distribution from the neuron’s linear model (see above) by setting the coefficients of side-of-object and its interactions to 0, and then simulating the neuron’s responses by adding random Gaussian noise with the variance equal to the residual variance of the model fit. The 95% confidence intervals of the null-hypothesis line were calculated by bootstrap, resampling scenes with replacement.

To calculate a consistency index we compiled the distribution of the neuron’s distribution of border ownership effects across scenes and deconvolved this distribution with the null-hypothesis distribution, using the DeconPdf function of the R decon library (Wang and Wang, 2011). The original distribution was first approximated with a Gaussian to eliminate aliasing artifacts.

#### Time course

The spike rate histograms were calculated for the population of cells that were border-ownership selective in both the standard square test and the natural scenes test. The border-ownership signal is the difference between the spike rate histograms for preferred and non-preferred side. The histograms were smoothed (Lowess, tension 0.12).

Latencies were determined by two-phase regression on the cumulative spike count histograms with 1 ms resolution of the population (Sugihara et al., 2011). For the overall responses, the spike counts were cumulated in the interval from 0 to 80 ms (*Spikes_pref_* + *Spikes_nonpref_*). For the border-ownership signals, the difference between the spike counts for preferred and non-preferred conditions were cumulated in the interval from 30 to 150 ms (*Spikes_pref_*-*Spikes_nonpref_*). The onset of the context effect was calculated using the cumulative differential spike counts in the full image conditions minus the cumulative differential spike counts in the patch conditions,

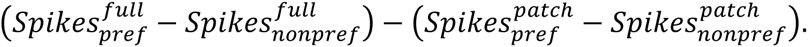

For the border-ownership signals the first leg of the regression fit was forced to zero, since the activity from the preferred and non-preferred conditions should cancel out before the onset of responses. In each case, the intersection of the regression lines defined the latency.

The standard deviations of the latencies were calculated by the bootstrap method (Efron and Tibshirani, 1993). The bootstrap was performed by resampling the stimulus presentations with replacement, within cell and condition. The number of presentations within cell and condition were kept the same as in the original data. For each resampled dataset, the latency estimates were calculated by two-phase regression as described above.

## Footnotes

1 Paolo Neo, www.public-domain-image.com/free-images/flora-plants/fruits/apple-pictures/red-apple-from-top

2 Two neurons responding to the same edge can have opposite directions of border ownership preference, and which direction we assign a positive value is arbitrary. Of course, the same assignment was used for all data from a given neuron.

